# Stomata: Gatekeepers of Uptake and Defense Priming by Green Leaf Volatiles in plants

**DOI:** 10.1101/2024.05.22.595386

**Authors:** Feizollah A. Maleki, Irmgard Seidl-Adams, Gary W. Felton, Mônica F. Kersch-Becker, James H. Tumlinson

**Affiliations:** Center of Chemical Ecology, Department of Entomology, The Pennsylvania State University, University Park, Pennsylvania 16802, USA

**Keywords:** E-3-hexen-ol, Defense priming, Fall army worm, Green leaf volatiles, Maize, Sesquiterpenes, Stomata, Z-3-hexen-ol

## Abstract

Plants adapt to balance growth-defense tradeoffs in response to both biotic and abiotic stresses. Green leaf volatiles (GLVs) are released after biotic and abiotic stresses and function as damage-associated signals in plants. Although, GLVs enter plants primarily through stomata, the role of stomatal regulation on the kinetics of GLVs uptake remain largely unknown. Here, we illustrate the effect of stomatal closure on the timing and magnitude of GLVs uptake. We closed stomata by either exposing plants to darkness or applying abscisic acid, a phytohormone that closes the stomata in light. Then, we exposed maize seedlings to *Z-3-hexen-1-ol* and compared the dynamic uptake of *Z-3-hexen-1-ol* under different stomatal conditions. Additionally, we used *E-3-hexen-1-ol*, an isomer of *Z-3-hexen-1-ol* not made by maize, to exclude the role of internal GLVs in our assays. We demonstrate closed stomata effectively prevent GLVs entry into exposed plants, even at high concentrations. Furthermore, our findings indicate that reduced GLV uptake impairs GLVs-driven induction of sesquiterpenes biosynthesis, a group of GLV-inducible secondary metabolites, with or without herbivory. These results elucidate how stomata regulate the perception of GLV signals, thereby dramatically changing the plant responses to herbivory, particularly under water stress or dark conditions.

We elucidate the role of stomata, small pores on plants’ leaf surface, in regulating the entry of green leaf volatiles, damage-associated signals, into plants, and thus influencing their signaling functions.

## INTRODUCTION

Plants inhabit dynamic environments that prompt them to develop different strategies to effectively balance growth and defense in response to both biotic and abiotic stresses (Huot *et al*., 2014). Plants have evolved mechanisms to mitigate the impacts of environmental stresses, such as changing their physical structure and chemistry (Taiz and Zeiger, 2015). As part of the chemical adaptation, plants release Green leaf volatiles (GLVs) from damaged tissues under thermal fluctuations, freezing conditions, variations in light and water availability, as well as mechanical damage, herbivory, and pathogen attacks (Ameye *et al*., 2017; Dudareva *et al*., 2013). GLVs diffuse through air or tissue and prepare neighboring plant cells for upcoming environmental stresses. Like other airborne signals, the delivery of GLVs depends on the stomatal aperture (Maleki, 2021; Aguirre *et al*., 2023; Aratani *et al*., 2023). Fast stomatal closure is one of the early physical responses of plants to environmental stresses, like water stress and wounding. Thus, the coincidence of this physical response with the release of GLVs as airborne signals poses a question about GLVs delivery to neighboring tissues. This becomes more critical if we consider the fragility of GLVs as small molecules and the chance of wind drift that restricts the radius of GLVs’ target site around the damage site (Schuman, 2023). While there is evidence of stomatal regulation over volatile organic compounds (VOC) emission and uptake (Aratani *et al*., 2023; Waterman *et al*., 2024), little is known about the magnitude of this regulation in the short term and with different exposure dosages.

Stomatal regulation (movement) is one of the major physical adaptations of terrestrial plants to survive different environmental stresses, and it impacts plants’ primary and secondary metabolisms. Stomatal closure occurs after environmental stresses, including flooding, elevated CO_2_ levels, heat, drought, high light, high salt, herbivory, and pathogen infection (Brodribb and McAdam, 2017). In this context, the activation of abscisic acid (ABA) represents one of the initial physiological processes that can function both locally and systemically to accelerate stomatal closure (Kuromori, *et al*., 2018). Closed stomata prevent the emission of H_2_O and intake of CO_2_, which both have dire physiological consequences, mainly turning down the photosynthesis rate and nutrient transport via xylem vessels (Taiz and Zeiger, 2015). Slowing down primary metabolism reduces substrates for some secondary metabolites. For instance, stomatal closure can reduce the biosynthesis of herbivore-induced sesquiterpenes, 15-carbon terpenoid compounds, perhaps by limiting the incorporation of CO_2_ into carbon backbones in maize (Block *et al*., 2017; Seidl-Adams *et al*., 2014). However, stomatal closure doesn’t always lead to a decline in secondary metabolite production. Instead, the plant’s response can be context-dependent. Under specific environmental stresses, plants may even ramp up production of targeted secondary metabolites to counter the stress. For instance, hyperosmotic stress in tea plants (*Camelia sinensis*) closes stomata and upregulates GLVs biosynthesis enzymes, and the accumulation of (*Z)-3-hexen-1-ol* (Z-3-HOL) inside leaves alleviates plant stress damage (Hu *et al*., 2020). After the change in abiotic factors like temperature, soil humidity, air humidity, and light, the emission of herbivore-induced VOCs is regulated through the stomatal aperture and the rate of VOC biosynthesis process (Gouinguené and Turlings, 2002; Niinemets *et al*., 2014; Niinemets and Reichstein, 2003).

The uptake of VOCs is regulated by the stomatal aperture and the concentration gradient between the inside and outside of plant leaves (Niinemets *et al*., 2014). The concentration gradient of VOCs varies based on reactivity and biosynthesis rate (Niinemets *et al*., 2014), physicochemical properties (Niinemets and Reichstein, 2003), and incorporation of absorbed compounds in the leaf metabolism (Matsui, 2016). Based on the physicochemical properties of VOCs, they can distribute (partition) in different leaf compartments, including lipid, aqueous, and gas phases. The partitioning between aqueous and gas phase, depicted as Henry’s law constant *(H)*, can explain the behavior of VOC molecules in the stomatal cavity (Niinemets and Reichstein, 2002). Some small VOC, like Z-3-HOL have low (<10) *H*, meaning their concentrations in the aqueous phase of the leaf is higher than gas phase. Whether the source of a soluble VOC aqueous pool is biosynthesis or uptake from the air, its lingering pool restricts the VOC uptake unless it is metabolized or converted to other derivatives (Tani *et al*. 2013). The conversion of up taken VOC to other metabolites and their conjugation reduces the pool of VOC inside the plant tissue and increases the concentration gradient between the inside and outside of the plants to uptake more VOC (Matsui, 2016). For example, the conversion of the GLV alcohol, Z-3-HOL to *Z-3-hexenyl acetate* (Z-3-HAC) controlled by acetyl-CoA:(Z)-3-hexen-1-ol acetyltransferase enzyme (D’Auria *et al*., 2007) happens minutes after plants uptake the external Z-3-HOL as has been shown by using ^13^CO_2_ in maize plants (Farag, *et al*., 2005). The esterification happens with similar rate for other GLV alcohols, like *(E)-3-hexen-1-ol* (E-3-HOL) an isomer not made by maize but can be converted to corresponding ester metabolite by maize (Yan and Wang, 2006). When stomata are open, the uptake of VOC depends on the diffusion gradient of the compounds in the water phase of leaf cells. However, when stomata are closed, there is a physical barrier to the uptake (Niinemets *et al*., 2014).

The airborne GLVs are delivered to adjacent plant tissues through stomata (Aratani *et al*., 2023) and induce a general response, such as Ca^2+^ burst (Aratani *et al*., 2023; Zebelo *et al*., 2012), followed often by jasmonic acid (JA) pathway activation in plants (Engelberth *et al*., 2004; Engelberth *et al*., 2013). Jasmonates are other lipid peroxidation products in plants induced by wounding or herbivore attacks and after exposure to GLVs (Arimura *et al*., 2008; Schmelz *et al*., 2003). For example, in maize, GLVs boost JA response and consequent volatile organic compound (VOC) emission (Engelberth *et al*., 2004). GLVs can trigger the induction of different classes of terpenoids and aromatic compounds in maize plants in a dose-dependent manner (Ruther and Fürstenau (2005). Recently, it has been shown stomatal closure dampens the intake of Z-3-HAC into maize and then decreases the induction of JA significantly and, consequently, terpenoid compounds (Aguirre *et al*., 2023).

GLVs activate the biosynthesis of terpenoids, attractants of noctuid larvae parasitoids (Kollner *et al*., 2008; Schnee *et al*., 2006), thus increasing the chance of successful indirect defense for maize. While the outcome of indirect defense is context-dependent (Turlings and Erb, 2018), GLVs’ direct impact on noctuid larvae has yielded contradictory results in maize. Some studies have reported GLVs biosynthesis pathway negatively affects *Spodoptera exigua* larvae performance (Christensen *et al*., 2013; Ton *et al*., 2007). In contrast, others found more attraction of *S. frugiperda* (fall army worm) and *S. littorlais* to GLVs (Carroll *et al*., 2006; Von Mérey *et al*., 2011, 2013). Such an attraction in the field conditions might lead to increased *S. frugiperda* (fall army worm) damage on the maize plants exposed to GLVs (Von Mérey *et al*., 2011). An interesting aspect of some noctuid armyworms (especially fall army worm) is their nocturnal feeding. As recent evidence suggests, stomatal closure can dampen signaling function of GLVs at intra (Maleki *et al*., 2024) and inter-plant (Aguirre *et al*., 2023) levels in maize, and the consequence of such regulation on the noctuid larvae begs more investigation.

Stomata are major pathways of GLVs entrance into the plant (Aguirre *et al*., 2023; Aratani *et al*., 2023; Waterman *et al*., 2024); however, the degree of stomatal regulation on the GLVs signaling has not yet been quantified. Thus, to fill this knowledge gap, we determined the role of stomata closure regulation on the uptake of the GLV alcohol Z-3-HOL and its subsequent induction and priming functions. To study the role of stomatal closure on the uptake of Z-3-HOL, we tested the following hypotheses: 1. Maize uptakes less Z-3-HOL after stomata closure; 2. Lower uptake of Z-3-HOL reduces plant stress response in terms of VOC production. 3. Pre-exposure to Z-3-HOL primes biosynthesis of other VOCs in maize plants after herbivory, but only if the stomata are open during exposure. 4. Less uptake of Z-3-HOL into plants leads to better performance of FAW larvae.

## MATERIALS AND METHODS

### Plants growing conditions

14-day-old Delprim maize seedlings were used for all experiments. Maize seeds were planted in a horticultural growing mix of Sungrow^®^ with ingredients; 70-80% Canadian Sphagnum peat moss, perlite, vermiculite, dolomite lime and wetting agent inside 6-inch pots. Nutrients were provided by mixing Osmocote^®^ pellets (15-9-12 NPK) with the soil before planting seeds. Seedlings were maintained in an environmental room with a daily temperature of 25 °C and a 16:8 hours Light to Dark regime. We used distilled water to irrigate plants every other day.

### Stomatal closure manipulations and general experimental methodology

To induce darkness, glass tubes containing excised seedlings were concealed under a thick white sheet. Since the darkness induces a transient release of GLV, after dark induction the volatiles were flushed by pushing air through the system without any filters attached at the top of the containers, and a recovery period of 2 hours was applied to prevent any confounding effect of GLV traces of dark shock on the exposure experiment. To close the stomata in the light, ABA was used at 250µM concentration dissolved in the feeding water of plant seedlings (Figure 1A). For this, 6.6 mg of ABA was dissolved in 100ml distilled water to make a master mix and 15 ml aliquot was poured in each 20 ml vial to trigger closure of stomata. A grace period of three hours was considered long enough to ensure the uptake of ABA by plants. Any potential side effects of ABA or darkness on the emission of VOCs were checked by running treated plants with ABA or darkness alone in parallel with control plants. To verify the closure of stomata during the experiments, weight of water vials containing cut plants were recorded before and after experiments. Less weight loss in the dark or after ABA treatment was interpreted as successful stomatal closure and lower transpiration (Supplementary Figure S1).

**Figure 1.**
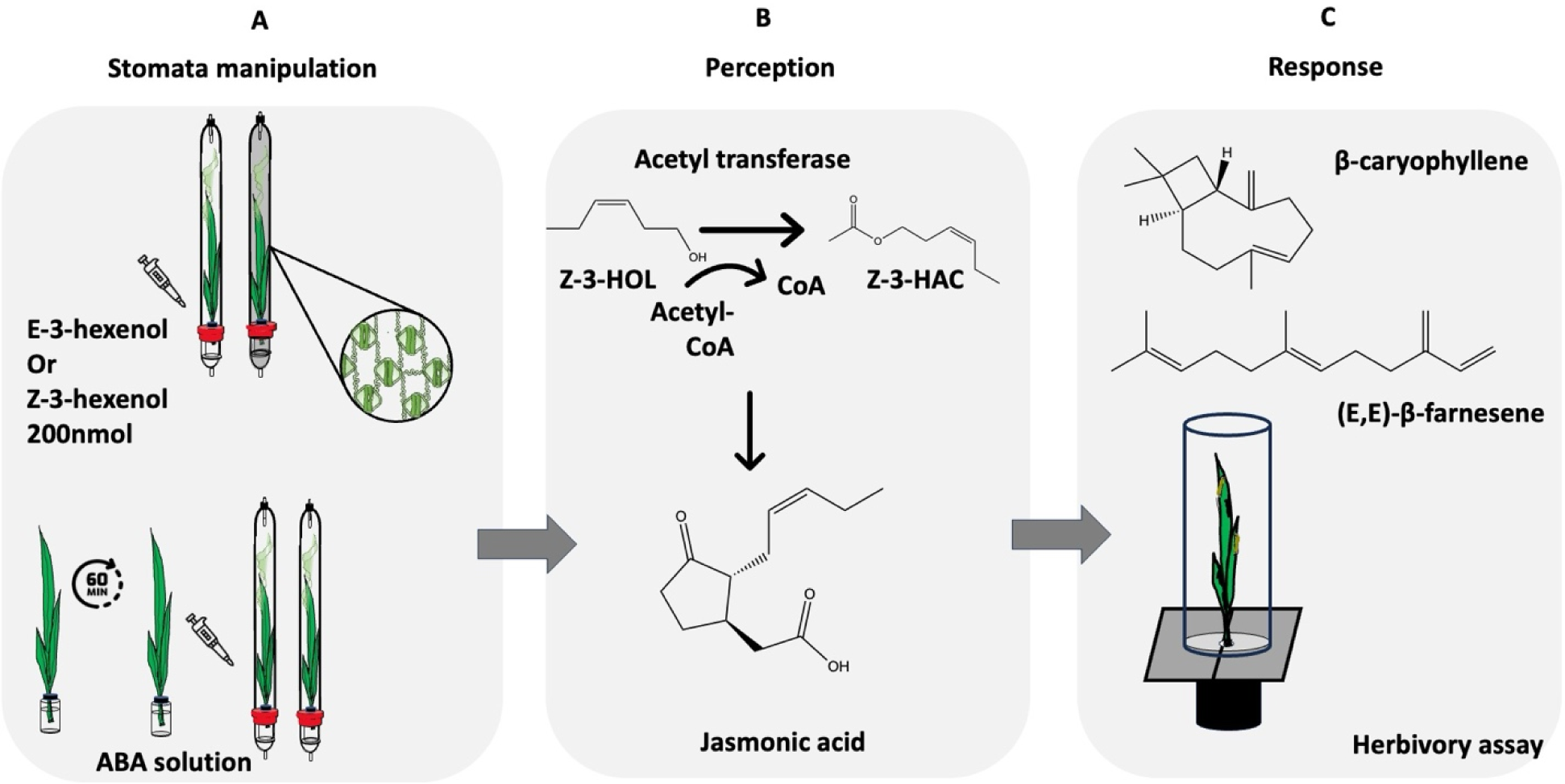
Experimental design and methodology used to test our hypothesis. A) Stomatal manipulation by darkness and Abscisic acid (ABA) application. B) We used the conversion of GLV alcohol to the related ester as a delivery index and measured free Jasmonic acid (JA) concentrations to convey the perception. C) The induction or priming effect of GLVs on sesquiterpenes biosynthesis was used as an estimate for the plant final response to GLV signals uptake. We also used a herbivory assay with FAW to address the direct consequences of stomatal regulation of GLV uptake on the defense response against FAW.

In addition, we used other methods to ensure stomatal closure after our manipulation techniques. We compared the stomatal aperture between treatments by measuring the length and width of stomatal openings on the glass slides of leaves nail polish. We brushed a little amount of nail polish on the leaf surface 1 hour after the start of the stomatal closure and removed the layer of nail polish after drying from the leaf surface. The nail polish layer was fixed under a glass cover on a glass slide, and then we took pictures of the slides at 20x and 40x magnification under the microscope which later were processed the microscopic picture with ImageJ software (National Institutes of Health) (Supplementary Figure S2). Furthermore, we used a porometer LI-COR® 600 (LI-COR Biosciences, Lincoln, Nebraska) to measure the stomatal conductance based on the stomatal conductance to water after darkness. We measured stomatal conductance by clipping the porometer to different spots on the leaves of multiple plants under each light conditions (Supplementary Figure S3).

The variety of timing and dosages in our experiments was based on different patterns of GLV release after herbivory (Engelberth *et al*., 2004; Turlings and Tumlinson, 1992) or studies on priming plant defenses with GLVs (Timilsena, *et al*., 2020; Waterman *et al*., 2024). Randomly selected seedlings were cut 1 cm above the first leaf and inserted in water through a cross-cut rubber septum cap in 20 ml vials. Cut seedlings were placed in 3-liter glass vessels with 0.5 L. min^-1^ charcoal-filtered air entering the vessels from the bottom, and the air was pulled at 0.5 L. min^-1^ from the top (Figure 1A). GLVs aliquots were pipetted on a cotton ball at the lower part of glass vessels. HayeSep-Q filters 80/100 (Supelco, Bellefonte, PA) were set on top of the vessels attached to pull air tubes. Depending on the purpose of the experiment, the timing of replacing filters and using dose(s) was different, as we describe below.

Dose-response assay in light vs dark: We quantified the uptake rate of Z-3-HOL (≥ 98%, Aldrich^®^) by maize after exposing 20 µl aliquots of a dilution series of 5, 2.5, 1.25, and 0.625 µg. µl^-1^ of Z-3-HOL (equivalent of 1000, 500, 250, and 125 nmol) in dichloromethane (DCM). Filters were replaced every 15 minutes. This experiment was conducted under light or dark conditions (Figure 2).

**Figure 2.**
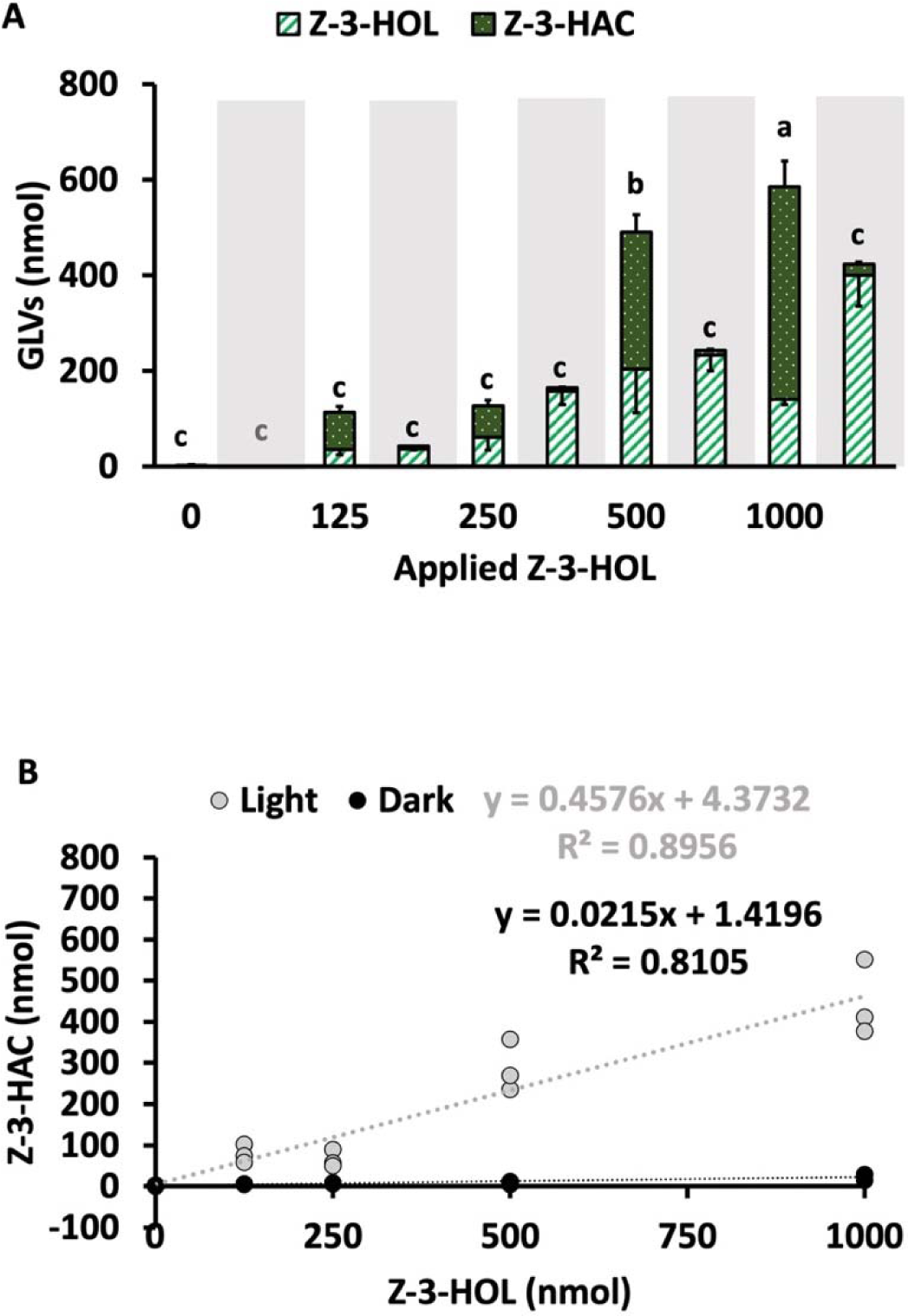
The conversion of different concentrations of Z-3-HOL to Z-3-HAC in light vs Dark-conditions by maize plants. A) The mass amount of headspace collected GLVs in light vs dark conditions after 30 min of plants exposure to different concentrations of Z-3-HOL dissolved in 20 µl of DCM. Aliquots of Z-3-HOL were pipetted on a small cotton piece at the start of exposure. B) The regression lines of dose-response relationship between applied Z-3-HOL and the amounts of Z-3-HAC. the mass amount of Z-3-HOL in nmol/container used for exposure experiment. Different letters indicate a significant difference between the amount of recovered Z-3-HAC at *P*<0.05. Error bars display standard errors (±SEM) (n=3).

Time course of GLV uptake in light vs dark: We recorded the time course of Z-3-HOL uptake in maize by using the highest concentration of Z-3-HOL in the previous experiment, 5 µg. µl^-1^ (1000 nmol), during 90 minutes in the light versus dark. We replaced filters every 5 minutes in the first 30 minutes and every 15 minutes for the last 60 minutes of the collection period. For each experiment, the recovery of GLVs from plants was compared to the corresponding value after pipetting GLV alcohol from empty glass vessels (Figure 3).

**Figure 3.**
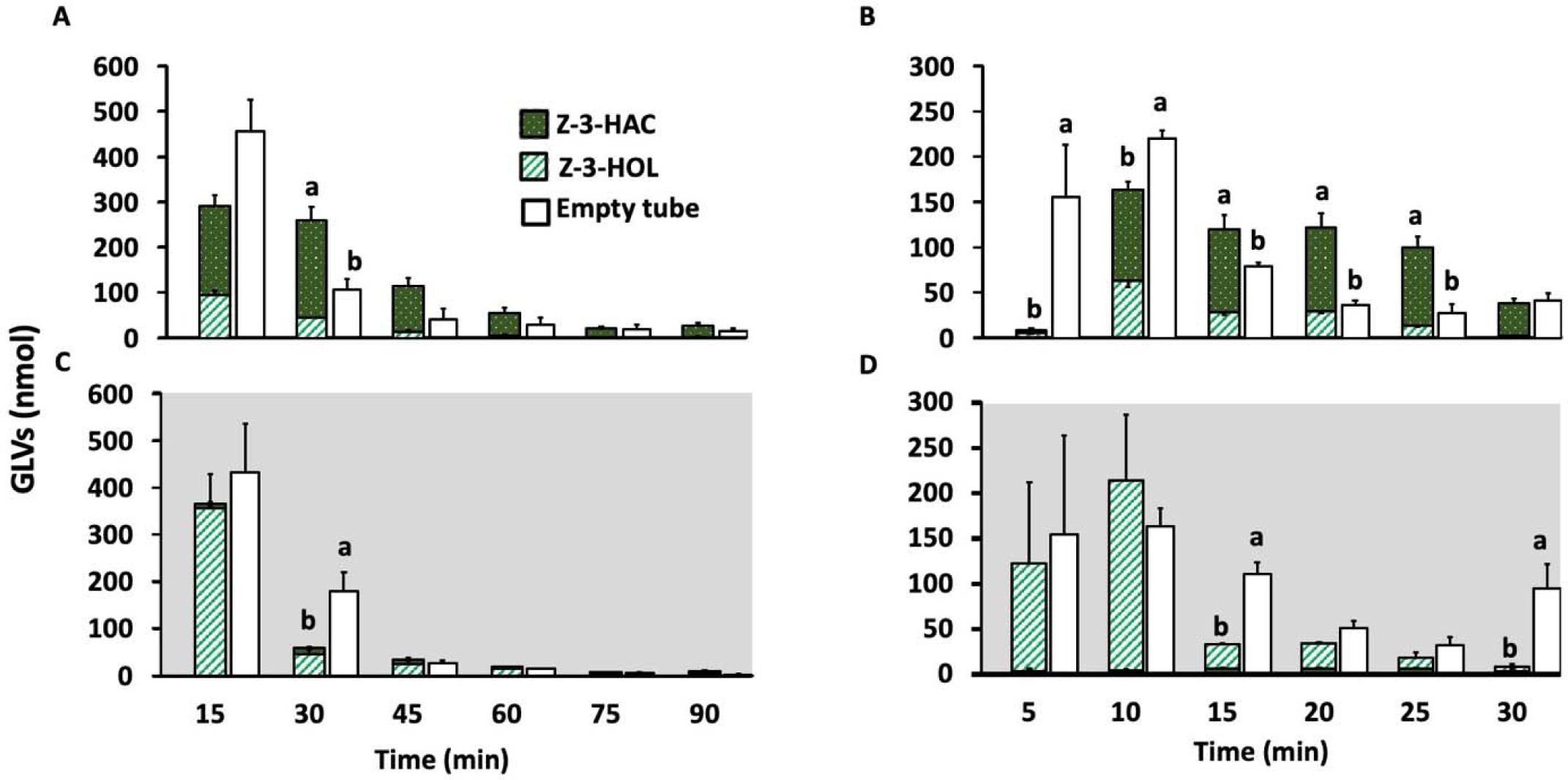
Time course of Z-3-HOL esterification by maize plants under light and dark conditions. A) GLVs headspace collections conducted over 90 minutes in different light conditions compared with the recovery of Z-3-HOL from empty tubes (white bar). B) Break down of the GLVs collected at 5-minute interval collections during the first 30 minutes in light conditions. C) GLVs headspace collections conducted over 90 minutes in dark conditions shown by a grey background. D) Break down of the GLVs collected at 5-minute interval collections during the first 30 minutes in the dark. Different letters indicate a significant difference between the sum of recovered GLVs from plants and recovered Z-3-HOL from empty tubes in each time point at *P<0.05*. Error bars indicate mean (±SEM) (n=3).

The effect of stomatal closure by darkness and ABA on the uptake of different GLV alcohols: To rule out the role of an internal pool of GLVs on the uptake of Z-3-HOL, we used E-3-HOL that is not made by maize in parallel to Z-3-HOL in every stomatal condition. Z-3-HOL and E-3-HOL (≥ 98%, Sigma Aldrich^®^) were used for experiments. A 20 µl aliquot of 1 µg. µl^-1^ concentration of Z-3-HOL or E-3-HOL in DCM was a physiologically relevant (Engelberth *et al*., 2004; Turlings and Tumlinson, 1992) for most assays. We pipetted 20 µl of a 1 µg. µl^-1^ concentration of GLV alcohols on the cotton ball at the bottom glass vessel and collected headspace volatiles for 1 hour, replacing filters every 15 min. The headspace assays were done in light vs dark or ABA vs non-ABA conditions (Figure 4).

**Figure 4.**
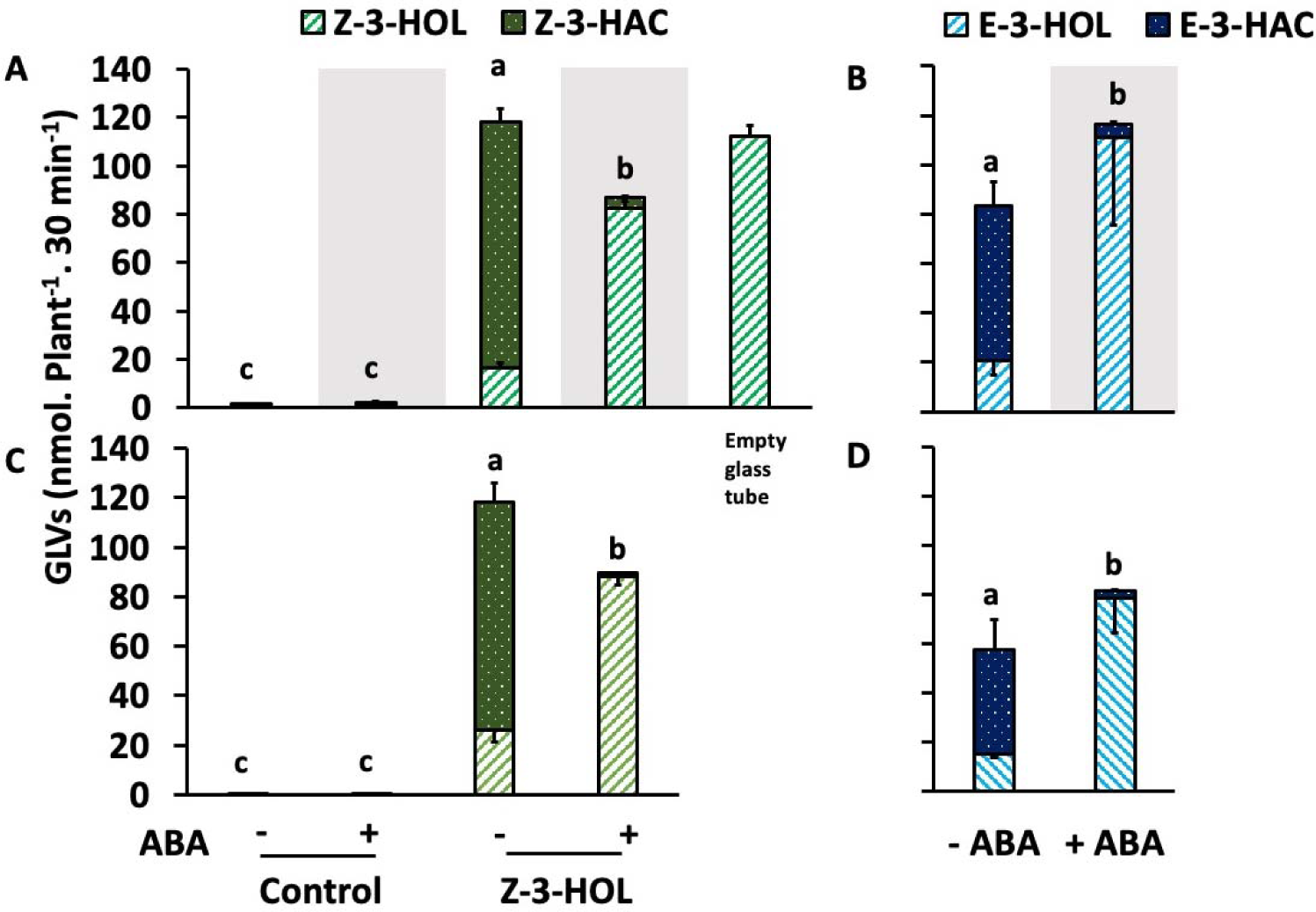
Assimilation of Z-3-HOL or E-3-HOL in under different stomatal aperture situations. (A) Collected GLVs from plants exposed to 200 nmol of external Z-3-HOL pipetted inside glass vessels after 30 min in the light vs dark conditions. The last bar on right, depicts the amount of Z-3-HOL from empty glass vessel without plants. B) Collected GLVs from plants exposed to 200 nmol of external E-3-HOL pipetted inside glass vessels after 30 min in the light vs dark conditions. C & D) Plants were treated with 250 µM ABA solution and then exposed to 200 nmol GLV alcohols, Z-3-HOL (C), E-3-HOL (D) as described before. Different letters indicate a significant difference between recovered acetate metabolites from plants under different light conditions or ABA application at *P<0.05*. Error bars display standard errors (±SEM) (n=3-5).

The role of stomatal closure on the uptake of Z-3-HOL during a pulsed exposure: To evaluate the effect of frequency of exposure on the uptake rate of GLV alcohol, we repeatedly pipetted 200 nmol of Z-3-HOL every hour for 5 hours inside the glass vessels. Maize stomata were manipulated as described before using darkness or ABA. After 5 hours of pulsed exposure, the dark-treated plants were exposed to light, and the ABA-treated plants were immersed in ddH2O to open the stomata after the exposure (Figure 5). After two consecutive 1-hour VOC collections, VOCs were collected overnight for 12 hours, as described before (Figure 5).

**Figure 5.**
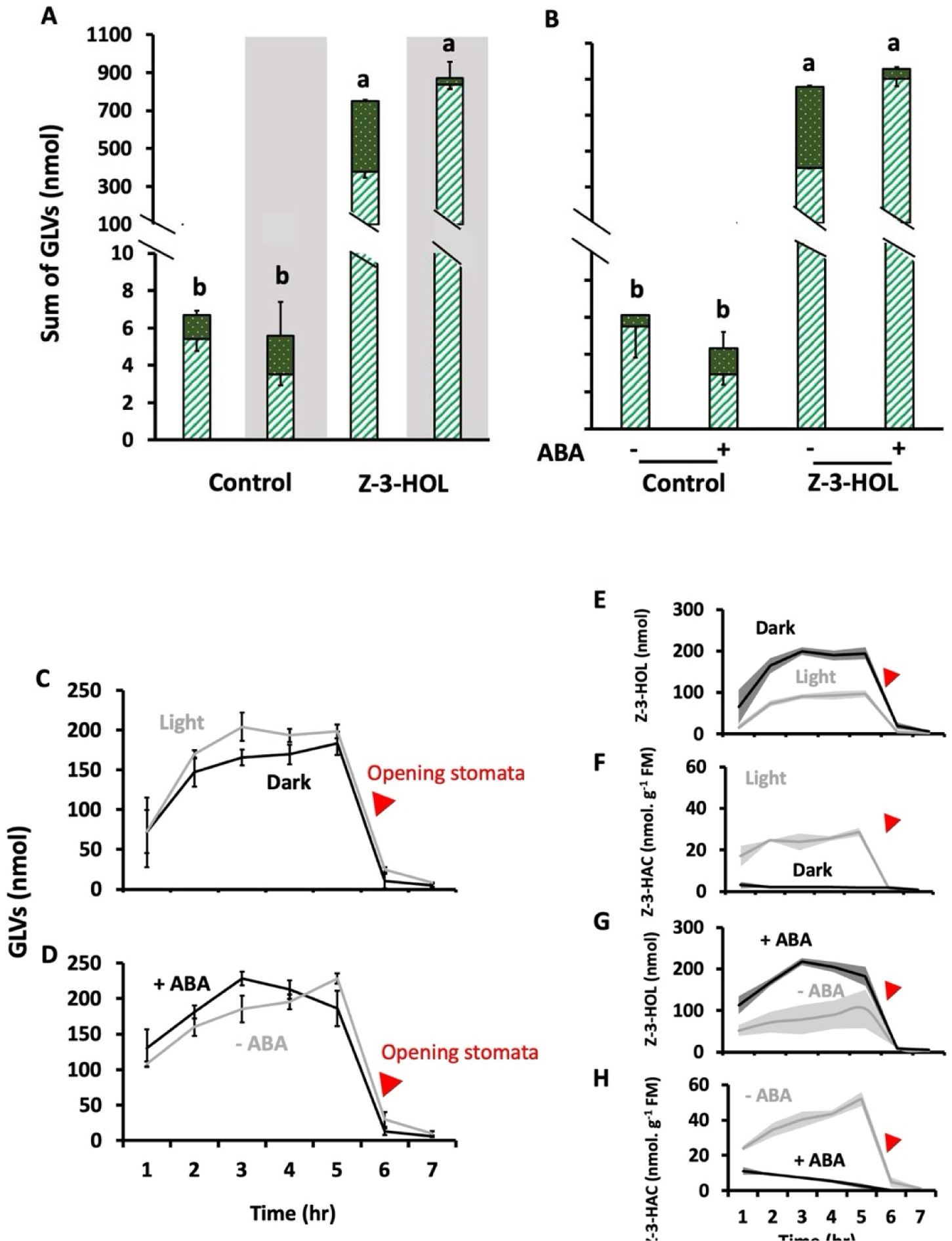
Tracking GLVs release from plants after exposure to multiple pulses of Z-3-HOL under different stomatal aperture conditions. Plants were exposed to 5 pulses of 200 nmol Z3HOL for 5 hours (200nmol/hr). After 5 hours of exposing plants with or without stomatal closure to Z-3-HOL, we opened the stomata (marked by red triangles) by removing the cover of the glass vessels (C) and removing plants from ABA solution and inserting them in the water (D). A) The sum of recovered GLVs from plants headspace after Z-3-HOL pulses in light vs dark. B) The sum of GLVs collected from plants headspace after Z-3-HOL pulses with or without ABA pre-application. C) The time course collection of GLVs from the plant’s headspace after the exposure to Z-3-HOL in light vs dark conditions over 7 hours and, D) after ABA application respectively. E) The time course of Z-3-HOL recovery from plants headspace in light vs dark conditions. The shadow black and grey show the standard errors of GLV quantifications in the dark, and light, respectively. F) The time course of Z-3-HAC recovery from plants headspace in light vs dark conditions normalized based on the fresh mass of plants. G) and H) The time course of Z-3-HOL recovery and Z-3-HAC from plants headspace with or without ABA application, respectively. Different letters indicate a significant difference between recovered acetate metabolites from plants under different light conditions or ABA application at *P<0.05*. Error bars display standard errors (±SEM) (n=3).

### JA measurement after plants exposure to Z-3-HOL in light vs dark

To test the effect of stomatal closure on the perception of Z-3-HOL in terms of JA induction, we exposed 14-day-old maize to 1µM Z-3-HOL in 7-L plexiglass cylinders under light or dark conditions. To induce darkness, plexiglass cylinders were covered by aluminum foil. After one 1-hour of recovery, we exposed plants to 1 µM Z-3-HOL (dissolved in DCM) pipetted on a cotton ball inside the cylinders. After 30 minutes, the time needed to detect JA induction in maize (Engelberth *et al*., 2004), all plants were harvested, fast frozen in liquid nitrogen, and stored in a -80°C freezer for further phytohormonal analysis. An adapted version of Schmelz *et al*., (2004) was used to quantify free JA. In brief, ground plant tissue was homogenized and transferred to DCM as solvent. Then the solvent evaporated, and the residue was dissolved in 1:9 methanol: diethyl ether and methylated by adding 2.0 M trimethylsilyl diazomethane in hexane. Methylation was stopped by adding 4 µl of hexane: acetic acid (880:120) to each sample. For more details, refer to the supplementary section. Methylated metabolites were collected on Hay-Sap Q filters according to Schmelz *et al*. (2003, 2004), and eluted with 150 μl of dichloromethane. Analyte detection was done using a quadrupole MS system (6890 GC coupled to a 5973 Mass Selective Detector; Agilent, Palo Alto, CA). The instrument conditions and quantification methods were as described in Engelberth *et al*. (2003) using selected-ion monitoring mode with retention times and [M+H]^+^ m/z ions are as follows: JA-ME (trans 12.28 min, cis 12.51 min, 225), Internal standards with retention times and [M+H]+ m/z ions were as follows: and dihydro-JA-ME (trans 12.34 min, cis 12.55 min, 227).

### Effect of stomatal closure on the priming defense response

To quantify the stomatal regulation over priming defense response, we expose plants to Z-3-HOL in the light versus dark conditions and then challenge them with FAW regurgitant. Plants were prepared and exposed following methods outlined in the previous section, except exposure to Z-3-HOL was 4 hours, and then all plants were challenged with FAW regurgitant in the light and inserted in new glass vessels for the rest of the experiment and an overnight collection for 12 hours. We used FAW regurgitant to mimic herbivory, as the plants simultaneously respond to the regurgitant like actual herbivory (Alborn *et al*., 1997; Signoretti *et al*., 2011). Fall army worm eggs were purchased (Benzon Research Inc., Carlisle, PA) and the larvae were reared on an artificial diet (Southland Products Inc., Lake Village, AR) under a 16:8 (L:D) cycle. We used 3^rd^-5^th^ -instar FAW larvae to collect regurgitant as described by Turlings *et al* (2000). Before collecting regurgitant, larvae were put on maize leaves for 24-48 hours before collecting regurgitant. For collecting regurgitant we held the back of caterpillar’s head and gently rub the mouth parts over a pipette pipe inserted through rubber septum screw cap inside a 4 ml glass vial over dry ice. A vacuum tube sucking air through a pipette tip passed through another hole on the rubber septum screw cap. Semi frozen collected regurgitant was centrifuged at 10000 rpm for 5 minutes and then the supernatant was stored in an Eppendorf tube in a -80 °C freezer for further use as elicitor. For elicitor application, 12 µl of regurgitant supernatant was mixed in 250 µl dd water and used as an elicitor to mimic herbivore attack.

### Herbivory bioassay

To evaluate the direct effect of stomatal regulation over Z-3-HOL priming, we used a dispenser that released 200 nmol. hr^-1^ of Z-3-HOL to expose the plants to Z-3-HOL. To make dispenser, we pipetted a 25 µl aliquot of mixture of Z-3-HOL and lanolin (1:1, V: V) paste inside a GC insert mounted in 2 ml GC vial. The suspension was done on a hot plate while glass wares were preheated to make a homogenized distribution of the Z-3-HOL inside the lanolin paste. After sealing the GC vial with a screw cap, a 3 µl glass pipette was inserted through the rubber septum such as 2 mm of the pipette tip remained out of the GC vial. 14-day-old plants were placed on an adjustable steel stand and were covered with plexiglass cylinders. Clean air entered the cylinders from the top with air flow 1 L. min^-1^ and a VOC collection system pulled air from each cylinder at 0.75 L. min^-1^. For dark treatment, the plexiglass cylinders were covered with aluminum foil at least 1 hour before the start of the exposure experiment.

After 5 hours of exposing plants to Z-3-HOL in either light or dark conditions, the dispensers were removed and 10 FAW neonates were placed on each plant. After 2 days of feeding, the survival rate and FAWs weight were measured. The next day, three larvae were chosen randomly (but with similar weight range among treatments) and were placed on the second set of Z-3-HOL exposed plants (as described above) for 5 days. At the end, survived FAW larvae were counted and weighted. To quantify leaf consumption, we scanned the leaves and later analyzed the images with ImageJ 2.0 software.

### Chemical analysis of VOC profile in the headspace of plants

HayeSep-Q filters were eluted with 150 μl of an elution buffer of hexane: dichloromethane (1:1, v/v) mixture containing 2 ng μl^-1^ 2-heptanone as an internal standard for GLVs and 4 ng μl^-1^ of nonyl-acetate as the internal standard for other VOCs. For quantification of GLV volatiles, a polar SPB-35 capillary column (29.3 m × 250µm × 0.25 µm film thickness; Supelco ®) was used housed in an Agilent model 6890 gas chromatograph equipped with a flame ionization detector. An aliquot of 1 µl was injected into the column using an automated injection system for the analysis. The program started with holding the oven temperature at 40 °C for 3 min. Then it increased the oven temperature at 8 °C min^-1^ to 100 °C, followed by another increase in the temperature at 5 °C min^-1^ to 200 °C and the final ramp at 40 °C min^-1^ to 260 °C which continued for 4 minutes. The Agilent Chemstation software was used to quantify all major compounds based on the detected intensity of a known quantity of 2-heptanone as the major internal standard.

The other VOCs were analyzed on a non-polar HP-5MS capillary column (30 m × 0.25 mm ID × 0.25 μm film thickness; Supelco, PA) installed in an HP model 6890 gas chromatograph equipped with a flame ionization detector. An automated injection system injected an aliquot of 1 μl per sample into the GC inlet. The initial GC oven temperature was held at 40 °C for 2 min, followed by a linear temperature increase of 4°C min^-1^ until 150° C was reached, after which the temperature increased at 40°C min^-1^ until 280°C. Then the oven was held at 280°C for 3 min. Agilent Chemstation software was used to calculate the peak area of detected volatiles. The quantity of volatiles was calculated relative to the peak area of the internal standard (here nonyl acetate). Most volatiles were identified based on spectrometric analysis of selected samples with an Agilent 6890N GC interfaced with an Agilent 5973N mass spectrometer (MS) detector and equipped with the same column. The method parameters used in GC-MS were similar to those used in GC-FID.

### Statistical analysis

Experiments were conducted either in a completely randomized block design, with dates of the experiments as blocking factors or in completely randomized design. The comparison uptake of GLV alcohol in plants with open or closed stomata consisted of four treatments and three-five replicates per treatment. Two-way analysis of variance (ANOVA) was used to test VOC emission changes in response to treatments; darkness or ABA and, exposure to GLV alcohol. In the case of non-significant blocking factors, data related to different dates was pooled and analyzed together. A Tukey HSD test was used to determine pairwise significant differences.

To analyze the relationship between the dose of Z-3-HOL and production of Z3HAC (light) and recovery of Z-3-HOL (dark) in the dose-response bioassay of Z-3-HOL in the light or dark, a linear regression was conducted, and then a one-way analysis of variance (one-way ANOVA) determined differences between the regression lines. For comparing the results of the GLV uptake time course assay, a student t-test was used. The collected Z-3-HOL or Z-3-HAC from plants in each interval was compared with the corresponding recovery of Z-3-HOL from empty glass vessels. To analyze the results of E-3-HOL volatile exposure of maize leaves, a student t-test was used to compare light versus dark treatment or ABA without ABA treatment.

Survival rate and leaf damage percentage among treatments were analyzed with the Chi-square test. A two-way ANOVA was used to detect the variation difference among treatments based on two factors of light condition and Z-3-HOL treatment for weight gain, and VOC emission. Then, significant results were followed by Tukey HSD. For long exposure and overnight VOC collection, peak area of VOCs were extracted and quantified by using a Python script (Timilsena, *et al*., 2020). Before analysis, the data was tested for homogeneity and normal distributions. In the case of non-normal variation, a box-cox transformation was done using Minitab 18 (State College, PA) software to normalize the data. Overnight VOC data were analyzed with Bray–Curtis dissimilarity metric using a permutational multivariate analysis of variance (PERMANOVA), with Z-3-HOL exposure and light condition as factors to map variation among treatments. We used R studio (R version 4.4.0; R core team, 2024) for PERMANOVA and related graphs.

## RESULTS

### Effect of concentrations on GLV uptake in different light conditions

In dark conditions, closed stomata controlled the flux of Z-3-HOL even at higher doses (Figure 2). In contrast, in light conditions, higher amounts of Z-3-HOL resulted in a higher conversion of the GLV alcohol to its acetate derivative. While we found a linear relationship between the applied dose of Z-3-HOL and Z-3-HAC in both conditions (Figure 2B), there was a significant difference between the recovered GLVs among treatments, indicating the impact of stomatal closure on the uptake of Z-3-HOL (Figure 2A). We found significantly higher Z-3-HOL recovery in the dark conditions based on the applied dose (*F_4, 22_*= 17.52, *P*= 0.000), light conditions (*F_1, 22_*= 9.90, *P*= 0.005) and a marginal difference based on the interaction of dose and light (*F_1, 22_*= 3.92, *P*= 0.017). The recovery of Z-3-HAC was significantly higher in the light conditions based on the applied concentrations of Z-3-HOL (*F_4, 20_*= 41.29, *P*= 0.000), light conditions (*F_1, 20_*= 153.24, *P*= 0.0), and the interaction of concentration and light conditions (*F_4, 20_*= 34.68, *P*= 0.000). Together, the results of these experiments suggest despite the increase in the Z-3-HOL concentrations, stomata can prevent the intake of Z-3-HOL and their impact is not dose dependent.

### Time course of Z-3-HOL esterification under different light conditions

A time course measurement of GLVs flux indicates maize plants assimilate most of the Z-3-HOL in the first 30 minutes after the start of the exposure (Figure 3A). Our statistical analysis shows significant differences between the sum of GLVs and recovered Z-3-HOL from empty vessels. In 0-15min; (*F_1, 5_* = 4.24, *P*= 0.109); 15-30 min: (*F_1, 5_* = 22.92, *P*= 0.009); 30-45 min: (*F_1, 5_* = 1.60, *P*= 0.274, box-cox transformed data); 45-60 min: (*F_1, 5_* = 0.07, P= 0.798); 60-75 min: (*F_1, 5_* = 0.10, *P*= 0.765, box-cox transformed data); 75-90 min: (*F_1, 5_* = 82.74, *P*= 0.485). In the empty tube (without plants), the highest amount of Z-3-HOL was collected in the first 15 minutes (white bars, Figure 3). However, the highest Z-3-HAC emission was recorded in the first 30 minutes for plants in the light (Figure 3A). In the first 30 min after the start of exposure, a similar release of Z-3-HAC happened in 15-minute intervals (Figure 3A), but in the first 5-minute interval, the assimilated GLV was very low (Figure 3B). We found significant differences between the sum of GLVs and recovered Z-3-HOL from empty vessels. In 0-5min; (*F_1, 5_* = 10.84, *P*= 0.005, box-cox transformed data); 5-10 min: (*F_1, 5_* = 56.01, *P*= 0.002); 10-15 min: (*F_1, 5_* = 11.58, *P*= 0.027); 15-20 min: (*F_1, 5_* = 34.89, *P*= 0.004); 20-25 min: (*F_1, 5_* = 32.22, *P*= 0.005); 25-30 min: (*F_1, 5_* = 82.74, *P*= 0.485). The first records of Z-3-HOL and Z-3-HAC were observed in the 2^nd^ 5 min interval, and this delay shows plants need time for the uptake and conversion of Z-3-HOL to Z-3-HAC (Figure 3B).

In the dark, the highest recovery of Z-3-HOL was achieved in the glass chambers during the first 30-minute interval (sum of six 5-minute intervals) with or without plant (Figure 3C). However, a significantly lower recovery was found for Z-3-HOL in plant-containing chambers during the second 15-minute interval (precisely in 3^rd^ and 6^th^ 5-minute intervals) In 0-5min; (*F_1, 5_* = 0.06, *P*= 0.822, box-cox transformed data); 5-10 min: (*F_1, 5_* = 0.41, *P*= 0.555); 10-15 min: (*F_1, 5_* = 34.14, *P*= 0.004); 15-20 min: (*F_1, 5_* = 5.93, *P*= 0.072); 20-25 min: (*F_1, 5_* = 2.4, *P*= 0.196); 25-30 min: (*F_1, 5_* = 10.6, *P*= 0.031) (Figure 3D).

### Assimilation of GLV alcohols under different light conditions and ABA application

Maize assimilated less Z-3-HOL and its isomer E-3-HOL after the closure of stomata in the dark. Z-3-HOL collected from the headspace of dark-exposed plants was higher than light-exposed plants, and respectively, Z-3-HAC, the conversion product of Z-3-HOL, was lower than in light-exposed plants (Figure 4A). After running statistical analysis (on box-coxed transformed data) on the recovery of Z-3-HOL from plants we found a significant effect of Z-3-HOL application, (*F_1, 21_*= 1156.06, *P* = 0.000; light, *F_1, 21_* = 107.80, *P* = 0.000, and their interaction *F_1, 21_* = 21.47, *P* = 0.000). For the recovery of Z-3-HAC the effect of treatment (*F_1, 21_* = 238.23, *P* = 0.000), light (*F_1, 21_* = 25.60 *P* = 0.000), and their interaction (*F _1, 21_* = 16.40, *P* = 0.001), were significant. The collected GLVs from light control plants were at basal level (≤ 1nmol), which shows that the internal pool of GLVs was low. Similarly, when using E-3-HOL, which is not produced by maize, higher amounts of E-3-HOL were recovered from the dark-treated plants, and higher E-3-HAC amounts were observed in the light-treated plants (Figure 4B). Our analysis shows a significant difference between light and dark-kept plants exposed to E-3-HOL for E-3-HAC (*F _1, 5_* = 35.73, *P* = 0.004, but not for E-3-HOL (*F _1, 5_* = 6.55, *P* = 0.063).

Our results confirm that maize assimilates less Z-3-HOL or E-3-HOL after ABA-induced stomatal closure in light-exposed plants. After stomatal closure, ABA-treated plants exhibited higher levels of Z-3-HOL, while plants without ABA application showed higher levels of Z-3-HAC (Figure 4C). A statistical analysis (on box-coxed transformed data) revealed a significant difference among treatments for recovered Z-3-HOL based on the Z-3-HOL application (*F _1, 11_* = 1087.9, *P* = 0.000), ABA (*F _1, 11_* = 19.82, *P* = 0.002) and their interaction (*F _1, 11_* = 34.52, *P* = 0.000). Also, for Z-3-HAC emission (box-coxed transformed data), the effect of Z-3-HOL treatment (*F _1, 11_* = 1844.78, *P* = 0.000), ABA (*F _1, 11_*= 287.16 *P* = 0.000), and their interaction (*F _1, 11_* = 573.14, *P* = 0.000) were significant. In addition, higher levels of E-3-HOL were recorded in ABA-treated plants compared to those without ABA application, and higher emission of E-3-HAC was recorded from ABA-free plants compared to ABA-treated plants (Figure 4D). Our analysis shows a significant difference between ABA and non-ABA treated plants exposed to E-3-HOL, in terms of recovered E-3-HOL, (*F_1, 11_* = 69.51, *P* = 0.000, box-coxed transformed data) and E-3-HAC (F*_1, 11_* = 57.29, *P* = 0.000, box-coxed transformed data).

Collectively, these results suggest the stomatal closure reduces the assimilation of GLV alcohols irrespective of light condition.

### The effect of stomatal aperture during the GLV exposure on the induction of VOCs

The pulsed exposure of plants to Z-3-HOL did not bypass closed stomata either in the dark (Figure 5 A), or with ABA application (Figure 5B), as the esterification results and the recovery of intact Z-3-HOL suggest. Despite similar sum of collected GLVs in all stomatal conditions (Figure 5A, and B), the composition of GLVs in headspace reveals most of Z-3-HOL enters plants in the light and without ABA and then converted to Z-3-HAC (Figure 5F, and H). The results analysis shows a significant difference among recovered Z-3-HOL treatments based on the exposure to Z-3-HOL (*F_1, 11_* = 931.06, *P* = 0.000), light conditions (*F_1, 11_*= 135.41, *P* = 0.000) and their interaction (*F_1, 11_* = 137.65, *P* = 0.000). In contrast, the emission of Z-3-HAC in the dark or after ABA treatment, remained low and steady throughout the experiment, showing the saturation status of plants due to the closure of stomata (Figure 5F, and H). We analyzed the box cox-transformed data and found significant difference among recovered Z-3-HAC from treatments based on the exposure to Z-3-HOL (*F_1, 11_* = 65.83, *P* = 0.000), light conditions (*F_1, 11_*= 6.35, *P* = 0.034), but not their interaction (*F_1, 11_* = 2.51, *P* = 0.15). These results correspond with higher recovery of Z-3-HOL that due to stomatal closure, remained intact and recovered through the assays (Figure 5E, G). Transitioning dark-treated plants to light allowed us to measure the possible pool of absorbed Z-3-HOL inside the plants. After exposing the dark-treated plants to the light, in the first two 1-hour intervals (6^th^ and 7^th^ hours on the graph), we collected more Z-3-HOL from pre-dark-exposed than light-exposed plants which might be leftover in the vessel or deposited on the surface of the leaves evaporated after the exposure to the light (Figure 5E). This trend continued, and more Z-3-HOL was recovered overnight from dark exposed plants to Z-3-HOL (Figure 6C, Supplementary Table S1). However, the amount of collected Z-3-HAC was significantly higher in light-exposed plants to Z-3-HOL (Supplementary Table S1). Thus, we demonstrate the stomatal closure not only restrict the uptake of GLVs after a high dose as a pulse (Figure 3) but also through a long period and with different pulses of Z-3-HOL (Figure 5).

**Figure 6.**
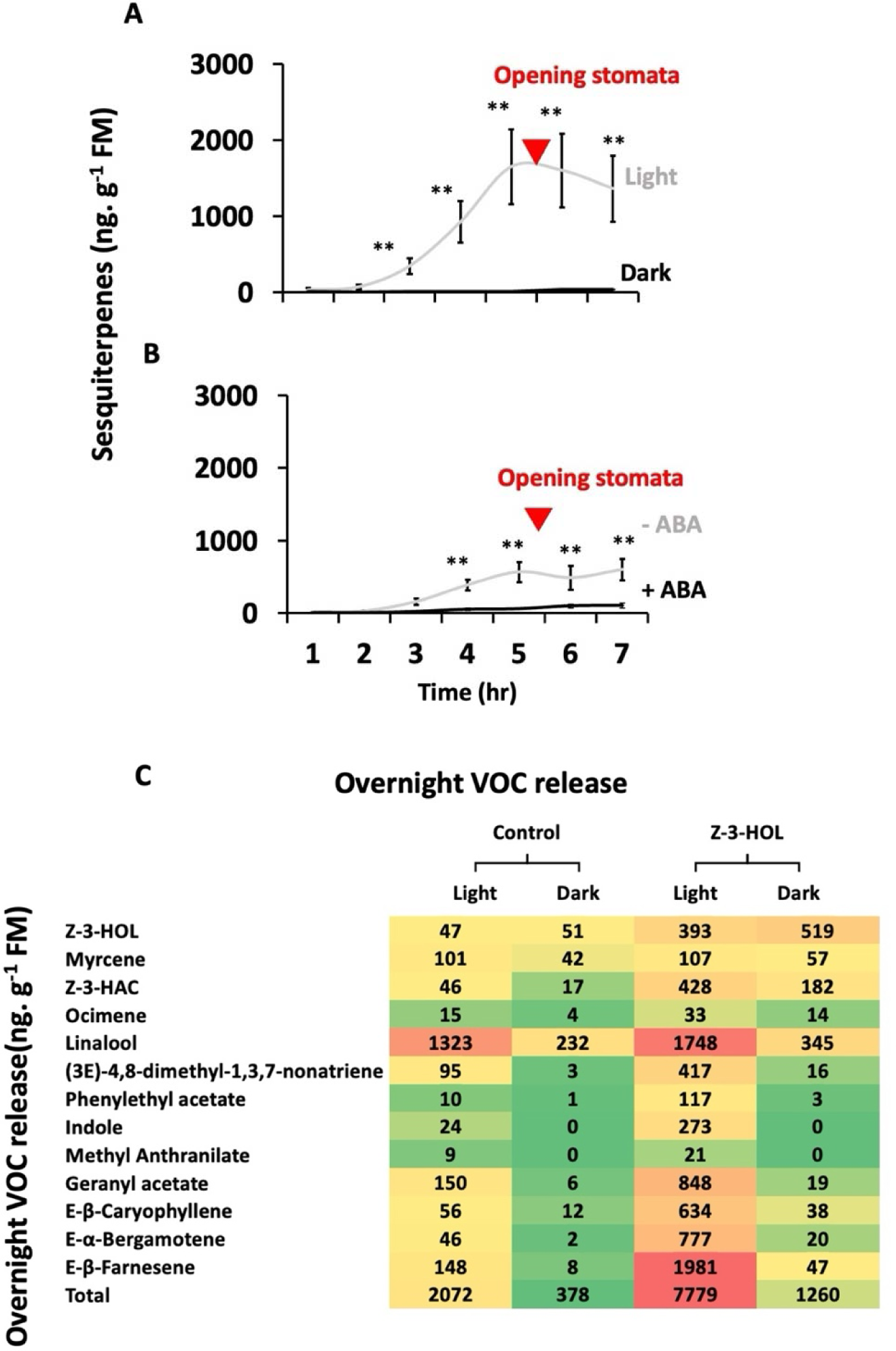
Time course of sesquiterpenes emission after exposing plants to Z-3-HOL pulses with stomatal closure during the exposure. As described in previous figure, we opened the plant stomata after 5 hours of Z-3-HOL exposure (marked with red triangle) and analyzed other VOCs besides GLVs over this period and overnight (12hrs). A) Time course of sesquiterpene emissions from the maize plants exposed to Z-3-HOL over 7 hours in light and dark conditions. B) Time course of sesquiterpene emission from maize plants with or without ABA application over 7 hours.). Error bars display standard errors (±SEM) (n=3) ** *P*≤ 0.01. C) Heatmap depicting the overnight (12 hours) mean VOC emission (ng. g^-1^ FM), lower amounts are shown in green, medium amounts in yellow and higher amounts in red.

We detected higher emission of terpenoid compounds after the exposure to Z-3-HOL in the light compared to when stomata were closed in the dark or after ABA application (Figure 6A, and B). Among the different terpenoids, the sesquiterpenes *β-caryophyllene*, *(E)-α-bergamotene* and *(E)-β-Farnesene* were the most representative in the headspace of maize plants (Figure 6A, and B). The high emission of sesquiterpenes continued even after plants were no longer exposed to Z-3-HOL and this effect lasted throughout the night (Figure 6C). The increase in the emission of sesquiterpenes started 2 hours after the beginning of the exposure and peaked after 5 hours. Plants exposed to Z-3-HOL in the light released more sesquiterpenes overnight (Figure 6C and Supplementary Table S2). Also running a PERMANOVA test on the overnight VOC profiles of plants revealed a significant effect of treatments (*F _3, 11_* = 18.80, *P* = 0.001, 999 permutations) (Supplementary Figure S5A).

### Jasmonic acid induction after exposing plants to Z-3-HOL in different light conditions

We quantified free jasmonic acid in the maize leaves during the peak uptake of Z-3-HOL, to assess the perception of Z-3-HOL with open or closed stomata. A two-way ANOVA (on box-coxed transformed data) revealed a significant difference among treatments for measured JA based on the Z-3-HOL application (*F _1, 11_* = 1087.9, *P* = 0.01), light (*F _1, 11_* = 19.82, *P* = 0.03) but, not for their interaction (*F _1, 11_* = 34.52, *P* = 0.083). Lower concentration of JA was found in response to Z-3-HOL in the dark, whereas the total free JA level was 50% more in plants exposed to Z-3-HOL in the light compared to the other treatments (Supplementary Figure S4).

### Stomatal aperture during the exposure determines the priming effect of Z-3-HOL

We manipulated Z-3-HOL intake to plants by closing stomata and found impaired priming function of Z-3-HOL after application of FAW regurgitant (Figure 7A). While we recorded an induction of sesquiterpenes during the Z-3-HOL exposure and a concomitant priming of such VOCs after herbivory challenge in the light, dark-exposed plants not only lack the induction phase but also the priming response (Figure 7A, B). Although FAW regurgitant triggered sesquiterpene emission in control plants in the light or dark conditions, the amount released was lower compared to plants exposed to Z-3-HOL in the light (Figure 7). The lowest emission of VOCs was observed in plants exposed to Z-3-HOL in the dark (Figure 7B, Supplementary table S2). The priming effect with FAW regurgitant and exposure to Z-3-HOL affected the emission of monoterpenes, homoterpenes, esters, aromatics, and sesquiterpenes. A PERMANOVA test found a significant effect of treatments on the VOC profiles of plants (*F _3, 11_* = 6.65, *P* = 0.006, 999 permutations) (Supplementary Figure S5B). This suggests the plant defense response was primed only when the Z-3-HOL could enter the plants in the light.

**Figure 7.**
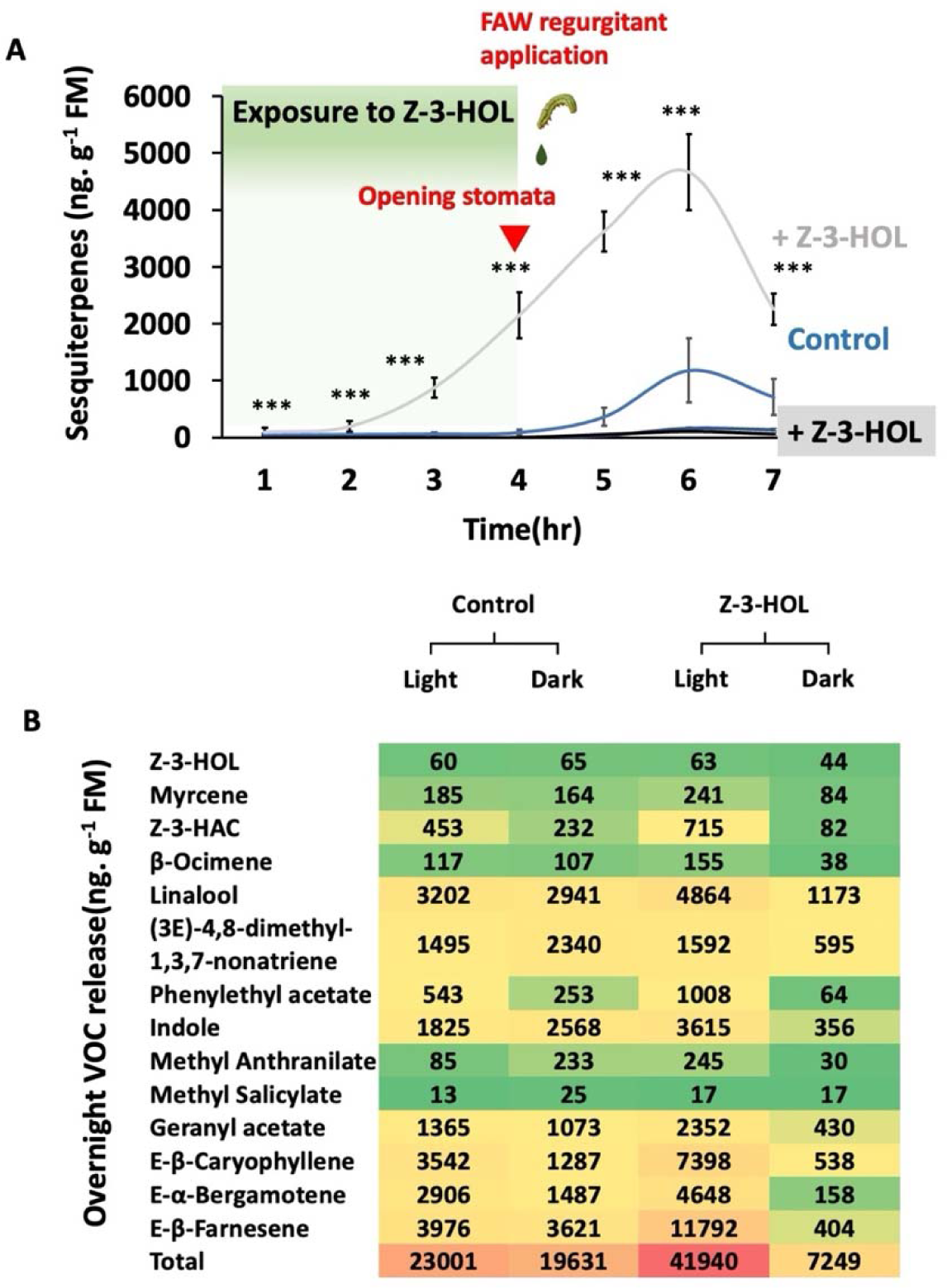
A) Time course of sesquiterpenes emission from plants headspace exposed to Z-3-HOL in light vs dark conditions (+Z-3-HOL) and then after application of FAW regurgitant. Cut seedlings were exposed to 4 pulses of 200nmol Z-3-HOL during a four-hour exposure (200nmol per hour) in light or dark conditions. After four hours all plants were moved to the light and challenged by 12 µl of FAW regurgitant in 250 µl of water. Error bars display standard errors (±SEM) (n=3) *** *P*≤ 0.001. B) A heatmap based on the average amount of each compound after 12 hours overnight VOC collection (ng. g^-1^ FM). lower amounts are shown in green, medium amounts in yellow and higher amounts in red.

### Dark kept plants suffered more from FAW feeding

After exposing plants to Z-3-HOL in different light conditions for 5 hours and then letting FAW larvae feed on plants, we found higher leaf consumption and weight gain in dark-treated plants regardless of Z-3-HOL treatment (Figure 8A, and B). We did not detect a significant effect of Z-3-HOL exposure on the weight gain of FAW larvae after running ANOVA two-way (*F_1,49_*= 1.33, *P*= 0.25, transformed data); however, FAW gained more weight when feeding on plants in the darkness, in both control and plants exposed to Z-3-HOL (*F_1,49_*= 4.21, *P*= 0.015, transformed data). It is noteworthy that we found a significant difference based on the interaction of block and Z-3-HOL treatment (*F_1,49_*= 9.97, *P*= 0.003, transformed data). FAW larvae consumed slightly more leaf area when feeding in dark-treated plants with a marginal significant difference (*F_1,22_*= 4.28, *P*= 0.053, transformed data), but Z-3-HOL exposure had no effect on FAW leaf consumption (*F*_1,22_= 0.22, *P*= 0.64, transformed data) (Figure 8B). After 3 days, the survival rate of FAW on control plants was 73.33% in light and 76.67% in dark conditions. In contrast, on Z-3HOL treated plants, the survival rates were 76.66% in light and 70% in dark conditions (Χ*^2^*= 0.478, df= 3, *P*= 0.924). Αnd after 7 days, the survival rate of FAW on control plants was 55.56% in the light and 88.89% in the dark conditions. On Z-3HOL treated plants, the survival rate was 72.22% in light or dark conditions (*Χ^2^*= 4.98, df= 3, *P*= 0.173).

**Figure 8.**
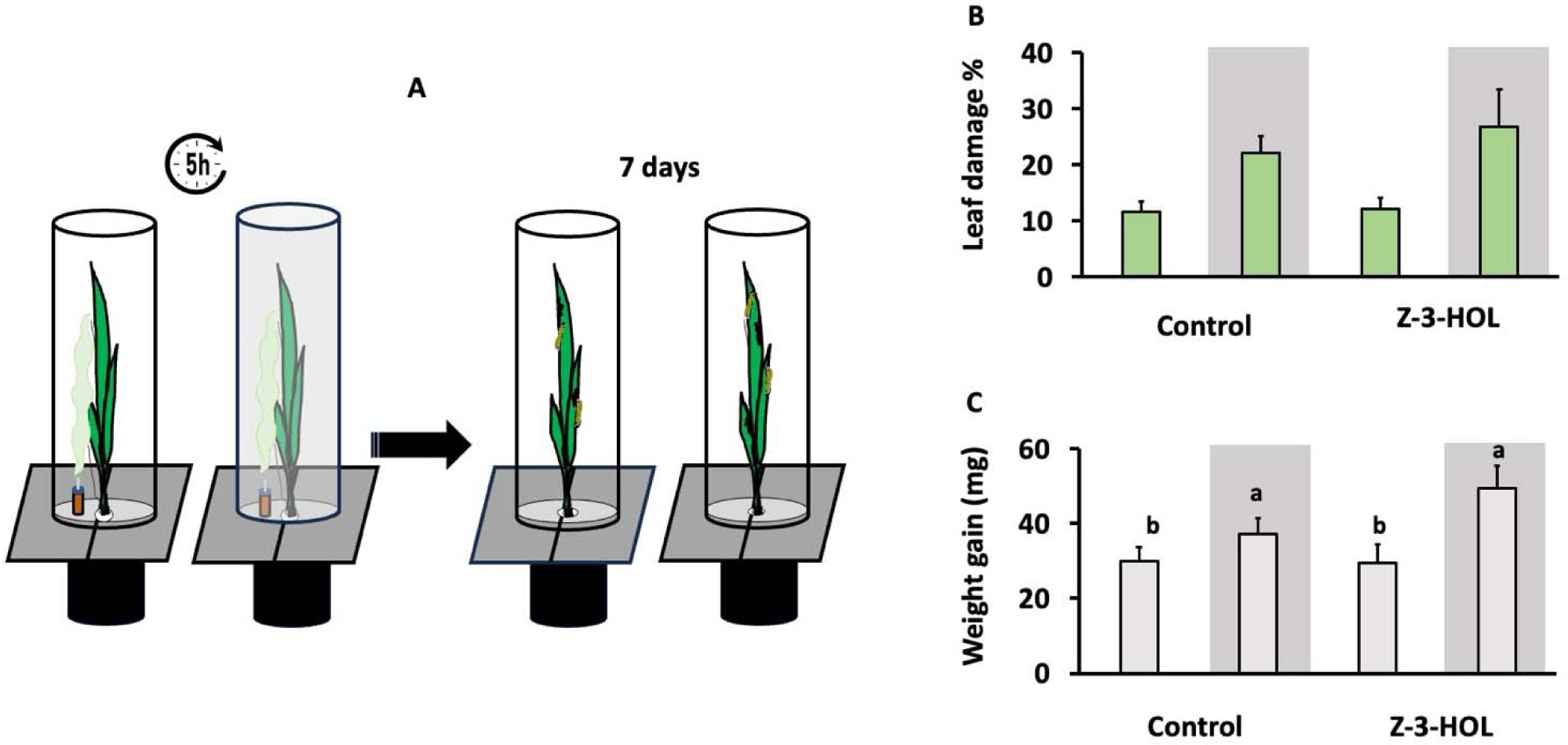
Fall armyworm (FAW) responses to light conditions and Z-3-HOL treatments. A) Plants were exposed to the lure of Z-3-HOL for 5 hours in the light vs dark conditions, and then were fed by FAW neonates for 7 days mass after 7 days. B) FAW leaf consumption in response to treatments. C) FAW larvae weight gain after 7 days. Different letters show significant difference based on light conditions. Error bars display standard errors (±SEM) (n=9-12).

## DISCUSSION

Stomata, regulators of water and CO_2_ exchange, impact plants’ physiology and primary metabolism; here, we highlight their critical role in tuning GLVs signaling as one of the early mechanisms after environmental stresses. We used different methods to manipulate stomatal closure to reduce GLV alcohol uptake, thus reducing the plant’s perception of these signals and, finally, terpenoids induction (Figure 6). Although there has been interest in the role of stomata in regulating GLV uptake (Aguirre *et al*., 2023; Aratani *et al*., 2023), our study demonstrates how stomatal aperture mediates the uptake of GLVs and the subsequent stress response of the plants. Despite closing stomata impairs the VOC-induction function of GLVs, indirect and direct effects on the herbivory are more nuanced, which we discuss in the following paragraphs.

Here we show that stomatal closure through two independent methods (darkness and ABA application) blocks the uptake of Z-3-HOL into maize leaves (Figure 4). Our results agree with what were reported about the role of stomatal regulation on the signaling functions of GLVs in the maize (Aguirre *et al*., 2023) and Arabidopsis (Aratani *et al*., 2023). Those studies relied majorly on phytohormones fluctuations (Aguirre *et al*., 2023) and Ca^2+^ bursts (Aratani *et al*., 2023) to show that stomatal closure impairs the GLVs perception in exposed plants. We used the conversion of GLV alcohols to esters (Farag, *et al*., 2005) as an index of delivery and illustrated stomatal closure prevents the entrance of GLV alcohols into plants and, thus, their conversion to GLV esters (Figures 2, 3, and 4). However, the biosynthesis of Z-3-HAC could also depend on the availability of acetyl-CoA and CHAT enzyme activity (D’Auria *et al*., 2007). Acetyl-CoA is independently synthesized in different subcellular compartments of plant cells, including cytosol, mitochondria, plastids, and peroxisomes (Fatland *et al*., 2002) and it has been shown that newly synthesized Acetyl-CoA can be incorporated in the turnover of Z-3-HOL to the acetate ester in cotton (Paré and Tumlinson, 1998) and maize (Farag, *et al*., 2005). Since darkness could reduce the acetyl-CoA pool (Hayashi and Satoh, 2008), we applied ABA to close stomata in the light condition, resulting in reduced conversion of Z-3-HOL to Z-3-HAC. It shows lower esterification of GLV alcohols is due to lower uptake rather than low acetyl-CoA pool (Figures 4C, D, and 5B, D, H). While the maize CHAT enzyme might have a different temporal activity than the Arabidopsis enzyme, the turnover of Z-3-HOL in the current study is very similar to previous studies (D’Auria *et al*., 2007) (Figures 3, 4).

Through both single (Figures 2, 3, 4) and pulsed exposure (Figure 5) of plants to GLV alcohols, we show that the effect of stomatal aperture on the uptake of GLVs is not time or dose-dependent. We detailed the stomatal regulation of Z-3-HOL uptake in the dark, which weakens the correlation between Z-3-HOL and the emission of Z-3-HAC at 30 minutes (Figure 3) and 5 hours (Figure 5). Stomatal closure in maize prevents the priming defense response of Z-3-HAC emitted from a slow-release rate dispenser (70 ng. h^-1^) overnight for 12 hours (Aguirre *et al*., 2023). Also, the early response of plants to GLVs, Ca^2+^ bursts (Aratani *et al*., 2023), and plasma depolarization (Zebelo *et al*., 2012) follow a dose-dependent pattern, which is delayed when stomata are closed by ABA application (Aratani *et al*., 2023). During herbivory, the fast diffusion of GLVs in the air leads to signal dilution and thus higher concentrations of GLVs (Aratani *et al*., 2023; Matsui *et al*., 2012) are only accessible in short distance meaning neighboring tissues (Schuman, 2023). Stomatal closure blocks the entrance of Z-3-HOL pulses (Figure 5), a mimic of frequent feeding and non-feeding phases of herbivory (Ji *et al*., 2017), and exhibits time is not a factor in the resistance of stomata. Our results strengthen the role of stomata in the early and long-term signaling responses of plants.

We demonstrate stomata regulate the uptake of E-3-HOL, a GLV that is not biosynthesized by maize, showing stomatal regulation of Z-3-HOL uptake is independent of the internal pool of Z-3-HOL (Figure 4). The absorbance of VOCs into plants depends on the chemical gradient of the compounds, which is determined by the concentration of VOCs in the aqueous phase of the plant tissue. Due to the low *H* value of GLVs, stomata regulate their emission and thus increase the half-life of their internal pool in plant tissues (Lin *et al*., 2021; Niinemets and Reichstein, 2003). In our system, cutting plant seedlings (Hatanaka, 1993; Matsui, 2006) and light-dark transition (Jardine *et al*., 2012) produced a burst of GLVs. We accounted for this by using E-3-HOL because maize plants convert E-3-HOL to E-3-HAC at a similar rate compared to Z-3-HOL (Yan and Wang, 2006) thus using E-3-HOL without an internal pool in maize refuted the effect of the GLV internal pool on our results (Figure 4). The similar structure of E-3-HOL and Z-3-HOL leads to the high affinity of CHAT enzyme for this GLV alcohol (D’Auria *et al*., 2007). If the internal pool of GLVs had contributed to the uptake of GLV alcohols, then the uptake rate should have been different for E-3-HOL compared to Z-3-HOL, which was not.

Our findings show the negative effect of stomatal closure on Z-3-HOL-induction of terpenoid and aromatic compounds (Figure 6), lasting even after reopening the stomata and herbivory challenge (Figure 7), which displays abiotic factors determine the signaling function of GLVs (Figure 6A, and B). Plants perceive Z-3-HOL significantly lower in the darkness (Figure 5A, C, E, and F), followed by lower terpenoid induction as a final response (Figure 6, and 7). While we only tracked free JA once in the dark exposed plants (Supplementary Figure S4), in another study, tracking JA did not show any induction after simulated herbivory when plants were exposed to GLV in dark conditions or after ABA application (Aguirre *et al*., 2023). Alternatively, it is also possible that plants assimilated less CO_2_ during the 6-hour darkness (or ABA treatment), lowering some VOCs biosynthesis. This can partially explain the VOC emission pattern as the sum of overnight VOCs in dark-exposed control plants is 5 times lower than control plants in the light (Figure 6C). Such a difference can be expressed more after herbivory for compounds like some terpenoids and aromatic compounds that are produced *de novo* upon herbivory damage (Paré and Tumlinson 1997). In maize, elevated CO_2_ lowers stomatal conductance, thereby decreasing the emission of *(E)-β-caryophyllene* and two homoterpenes, *(3E)-4,8-dimethyl-1,3,7-nonatriene* and *(3E,7E)-4,8,12-trimethyl-1,3,7,11-tridecatetraene* after application of *S. exigua*’s oral secretion (Block *et al.,* 2017). Also Seidl-Adams and colleagues in (2014) reported plants accumulate *E-B-farnesene* if stomata were closed by darkness or ABA after challenging plants with a fatty acid conjugate elicitor. In our experiments, reopening stomata after the GLV-exposure period was not followed by terpenoid induction or emission from potent storage in the tissue (Figure 6A, and B), consistent with our previous study (Maleki *et al*., 2024). While we found that plants perceive GLV alcohol as an imminent danger signal and increase terpenoids biosynthesis at neighboring tissue, this perception is not activated in the dark (Figure 6, and Supplementary Table S2) or in the plants treated with ABA (Aguirre *et al*., 2023).

The physical environment limits plants’ resources to spend on expensive secondary metabolites under herbivory (Herms and Mattson, 1992; Stamp, 2008), and our results can be a new dimension in growth-defense balance with complex outcomes for plants and herbivores. Different patterns of overnight VOC emission after Z-3-HOL exposure under light or dark conditions and with or without herbivory challenge depict part of this complexity (Supplementary Figure S5A, and B). Our results suggest stomatal closure during exposure to GLVs reduces their impact in terpenoids induction and eventually in natural enemies’ attraction (Schnee *et al*., 2006), as indirect defense measures to the damaged leaves, which can impose lower pressure on herbivores (Kersch-Becker, *et al*., 2017). In addition to darkness and ABA-induced stomatal closure, herbivory can cause stomatal closure due to wounding and some effectors in herbivores’ saliva (like glucose oxidase), (Lin *et al*., 2021) highlighting the fitness benefits of stomatal manipulation for herbivores. On the other hand, closing stomata protects plants against harsh environmental conditions like drought, flooding and is a conservative feature of plants adaptation to live on earth (Hetherington and Woodward, 2003). Thus, the overall outcome of herbivory in such situations on plants can be very context-dependent, as, for example, plants under overnight herbivory differ vastly from plants under drought.

Whereas our results suggest stomatal closure can regulate plants’ indirect defense driven by GLVs (Figures 6 and 7), we found no effect of such regulation on the FAW performance (Figure 8). While we selected FAW based on their overnight feeding (Capinera, 2001; Hardke, *et al*., 2015), the tolerance and suppressing maize defense responses are reported for FAW larvae as specialist herbivores (Glauser *et al*., 2011; Köhler *et al*., 2015; Maag *et al*., 2014). In addition, Spodopteran larvae’s attraction to GLVs has been reported in the lab (Mérey *et al*., 2013), and field settings (Mérey *et al*., 2011), presumably due to GLVs feeding stimulation activity (Mérey *et al*., 2013). Our FAW bioassays show marginal differences based on light conditions (Figure 8), meaning FAW gained more mass (Figure 8A) and fed more (Figure 8B) on dark-exposed plants regardless of Z-3-HOL application. Perhaps using a more generalist armyworm species (like beet armyworm) *S. exigua* (Ton *et al*., 2007), with lower resistance to maize defense traits, would lead to different results and could be a good follow-up study.

Our study underscores the critical role of stomata in GLVs perception by blocking the entry of these airborne signals and thus interrupting their function in terms of inducing or priming indirect defense responses. In the dilemma of growth or defense, stomata, as physical adaptation of plants to abiotic stresses regulate defense chemistry. This regulation happens at the early steps of signal processing (delivery) to save plants energy for coping with abiotic stress or defense response at the right time. The low defense response in these situations could have neutral or advantageous outcomes for herbivores, which is worth further study.

## Supplementary data

The following supplementary data are available at JXB online.

Figure S1. Measuring plants water uptake after ABA (A) and darkness (B) exposure to verify stomatal closure.

Figure S2. Demonstration of stomatal closure based on the measurements of width and length of stomatal opening after 2 hours of (A) darkness and ABA application (B).

Figure S3. Impact of darkness on the stomatal conductance (gsw)

Figure S4. Jasmonic acid measurement in maize after exposure to Z-3-HOL in light vs dark conditions. Figure S5. Multivariate analysis of plants’ overnight volatile organic compounds profiles after exposure to Z-3-HOL under different light conditions without (A) and with (B) herbivory challenge

Table S1. Overnight volatile compounds emitted by maize plants after exposure to Z-3-HOL in the light versus dark conditions.

Table S2. Overnight volatile compounds emitted by maize plants after exposure to Z-3-HOL in the light versus dark conditions and FAW regurgitant application.

## Acknowledgment

We appreciate Nathaniel McCartney’s mentorship in gas chromatography analysis and previous members of the late Dr. Tumlinson lab for their feedback during the research. We are thankful to Danilo Ferreira for helping us in running R codes related to PERMANOVA. This paper is dedicated to the memory of Dr. Tumlinson who supported the project until the end.

## Notes

### Competing Interest Statement

The authors have declared no competing interest.

